# Novel insights into karyotype evolution and whole genome duplications in legumes

**DOI:** 10.1101/099044

**Authors:** Melissa M.L. Wong, René E. Vaillancourt, Jules S. Freeman, Corey J. Hudson, Freek T. Bakker, Charles H. Cannon, Wickneswari Ratnam

**Affiliations:** School of Environment and Natural Resource Sciences, Faculty of Science and Technology, Universiti Kebangsaan Malaysia, Malaysia; School of Biological Sciences, University of Tasmania, PB 55, Tasmania, Australia; Tasmanian Alkaloids, Australia; Biosystematics Group, Wageningen University, Wageningen, The Netherlands; Center for Tree Science, The Morton Arboretum, USA

## Abstract

Legumes (family Fabaceae) are globally important crops due to their nitrogen fixing ability. Papilionoideae, the best-studied subfamily, have undergone a Whole Genome Duplication (WGD) around 59 million years ago. Recent study found varying WGD ages in subfamilies Mimosoideae and Caesalpinioideae and proposed multiple occurrences of WGD across the family based on gene duplication patterns. Despite that, the genome evolution of legume ancestor into modern legumes after the WGD is not well-understood. We aimed to study genome evolution at the subfamily level using gene-based linkage maps for *Acacia auriculiformis* and *A. mangium* (Mimosoideae) and we discovered evidence for a WGD event in *Acacia*. In additional to synonymous substitution rate (Ks) analysis, we used ancestral karyotype prediction to further corroborate this WGD and elucidate underlying mechanisms of karyotype evolution in Fabaceae. Using publicly available transcriptome resources from 25 species across the family Fabaceae and 2 species from order Fabales, we found that the variations in WGD ages highly correlate (R=0.8606, p-value<0.00001) with the divergence age of *Vitis vinifera* as an outgroup. If the variation of Ks is corrected, the age of WGDs of the family Fabaceae should be the same and therefore, parsimony would favour a single WGD near the base of Fabaceae over multiple independent WGDs across Fabaceae. In addition, we demonstrated that genome comparison of Papilionoideae with other subfamily provide important insights in understanding genome evolution in legumes.

## INTRODUCTION

The legumes (family Fabaceae) are important crops worldwide due to their unique ability to fix nitrogen in the soil through interaction with microbes. Fabaceae is traditionally classified into three subfamilies, namely Papilionoideae, Mimosoideae and Caesalpinioideae, (Polhill, 1994), however, a new classification is currently underway in order to better reflect phylogenetic relationships (The Legume Phylogeny Working Group 1, 2013). Molecular phylogenetic analyses suggest a paraphyletic Caesalpinioideae comprises the earliest diverging lineages with Mimosoideae and Papilionoideae forming clades within it (Cardoso et al., 2012). The interval between origin and diversification of Fabaceae was estimated to be very short (about 1-2.5 million years) and Papilionoideae is thought to have diverged from Mimosoideae and Caesalpinioideae about 60 Million years ago (Mya) (Lavin et al., 2005).

The availability of plant genome sequences provides evidence that all plants are paleopolyploids as they have undergone several rounds of Whole Genome Duplication (WGD) followed by diploidization (Blanc and Wolfe, 2004; Eckardt, 2004; Schlueter et al., 2004; Tang et al., 2008; Jiao et al., 2011; Vanneste et al., 2014). Detecting these ancient WGDs is complicated by the occurrence of genomic rearrangement, gene loss, gene gain and transposable elements which often change gene content and order. All eudicots including legumes have undergone at least three rounds of WGD which have long been considered as an evolutionary force behind the origin and diversification of angiosperms (De Bodt et al., 2005; Jiao et al., 2011; Schranz et al., 2012). Early study based on examination of chromosome number found that most legumes are fundamentally tetraploids and predicted seven pairs of chromosomes as the ancestral chromosome set of legumes (Goldblatt, 1981). Most genomic research in legumes has focused on Papilionoideae due to their economic importance and large members including six published genome sequences[*Medicago truncatula* (Cannon et al., 2006), *Lotus japonicus* (Sato et al., 2008), *Glycine max* (Schmutz et al., 2010), *Cajanus cajan* (Varshney et al., 2011), *Cicer arietinum* (Jain et al., 2013; Varshney et al., 2013) and *Vigna radiata* (Kang et al., 2014)]. Genome comparisons have showed conserved syntenic blocks between papilionoid genomes with higher loss of synteny as phylogenetic distance increases (Young and Bharti, 2012). Previous studies based on comparative genomics analysis have suggested that one WGD event dated around 58-59 Mya occurred in Papilionoideae (Blanc and Wolfe, 2004; Schlueter et al., 2004; Pfeil et al., 2005; Cannon et al., 2006; Bertioli et al., 2009) with an additional one occurring approximately 13 Mya, exclusively in genus *Glycine* (Schmutz et al., 2010). The Papilionoideae WGD is thought to have played an important role in rapid diversification and origin of novel traits such as nodulation (Soltis et al., 2009; Young and Bharti, 2012).

In comparison with the largest and well-studied Papilionoideae, little is known about the genome organization and evolution of Mimosoideae and Caesalpinioideae, which contain many species of economic and ecological interest (such as mimosoids *Acacia, Prosopsis* and *Mimosa* and caesalpinioids *Caesalpinia, Senna* and *Tamarindus*). In these two subfamilies, relatively few genomic resources were available for comparative genomic analysis until recently. The use of Next Generation Sequencing has generated large amount of DNA sequence data from transcriptomes of the caesalpinioid *Chamaecrista fasciculate* and *Eperua falcata*(Brousseau et al., 2014), and the mimosoids *Acacia harpophylla, A. auriculiformis, A. mangium* and *Prosopsis alba* (Cannon et al., 2010; Lepais and Bacles, 2011; Wong et al., 2011). A study based on phylogenetic analyses of gene families and synonymous substitution rate (Ks) analysis found no evidence for WGD in *Chamaecrista fasciculata* (Cannon et al., 2010) and hinted that the Papilionoideae WGD has not occurred in Mimosoideae and Caesalpinioideae. However, a follow up study that sampled 20 species across the family and 17 outgroup species suggested multiple occurrence of WGD within and near the base of Fabaceae (Cannon et al., 2015). To further elucidate genome evolution and diversification of the Fabaceae, comparative genomic analyses needs to involve a broader range of taxa and use a variety of analytical approaches.

We aimed to understand Fabaceae genome evolution using new evidence from representatives of Mimosoideae and Caesalpinioideae and improved information from model legumes. The initial objective of this study was to investigate the genome structure and organization of *Acacia* in comparison to *M. truncatula* to facilitate the transfer of knowledge into *Acacia* genomic-assisted breeding programs. To provide novel insights into genomes of Mimosoideae, we construct linkage maps for *A. auriculiformis* and *A. mangium*. We compared them to the modern genome of *M. truncatula* and discovered a WGD in *Acacia* based on syntenic patterns. In an attempt to validate this WGD event, we performed Ks analysis in 29 species from 24 taxa from the three Fabaceae subfamilies and 2 taxa from the order Fabales. In addition, we predicted the ancestral genomic blocks from *M. truncatula* and tracked their positions in the modern genome of *M. truncatula* and *Acacia* to elucidate the underlying mechanisms of karyotype evolution in Fabaceae.

## RESULTS

### Comparison of the *M. truncatula* chromosomes with *A. auriculiformis and A. mangium* linkage maps

We constructed linkage maps for *A. auriculiformis* and *A. mangium* (subsequently referred to collectively as *Acacia*) using an *A. auriculiformis* × *A. mangium* F_1_ pedigree and compared each *Acacia* linkage map with the *M. truncatula* genome to assess gene synteny and colinearity (See Supplementary Figure 1 online). A total of 239 and 163 markers were mapped to fourteen linkage groups (LGs) in *A. mangium* and *A. auriculiformis*, respectively(Supplementary Table 1 online). Thirteen LGs, equal to the haploid number of chromosomes (n=13), were identified based on 54 common markers in both maps. We observed no major genomic rearrangement between the two *Acacia* genomes. About 92.5% and 92% of markers mapped in *A. mangium* and *A. auriculiformis* and respectively contained orthologous *M. truncatula* genes which formed 38 and 46 syntenic blocks respectively with *M. truncatula* genome (Figure 1A and 1B; Supplementary Table 1 online). Twenty four syntenic blocks (63%) from *A. auriculiformis* were found in *M. truncatula* genomic regions that showed overlap with syntenic blocks from *A. mangium,* despite the lower marker content and density. Most *Acacia* LGs were syntenic to two to three *M. truncatula* chromosomes, involving an entire *M. truncatula* chromosome arm in *A. auriculiformis* LG2, and *A. mangium* LG2 and LG6. Interestingly, we found three *Acacia* LGs (LG2, 9 and 11) to be syntenic to *M. truncatula* chromosome (Mt) 6 and these blocks covered the TIR-NBS-LRR megaclusters which are important for disease resistance.

**Figure 1.**
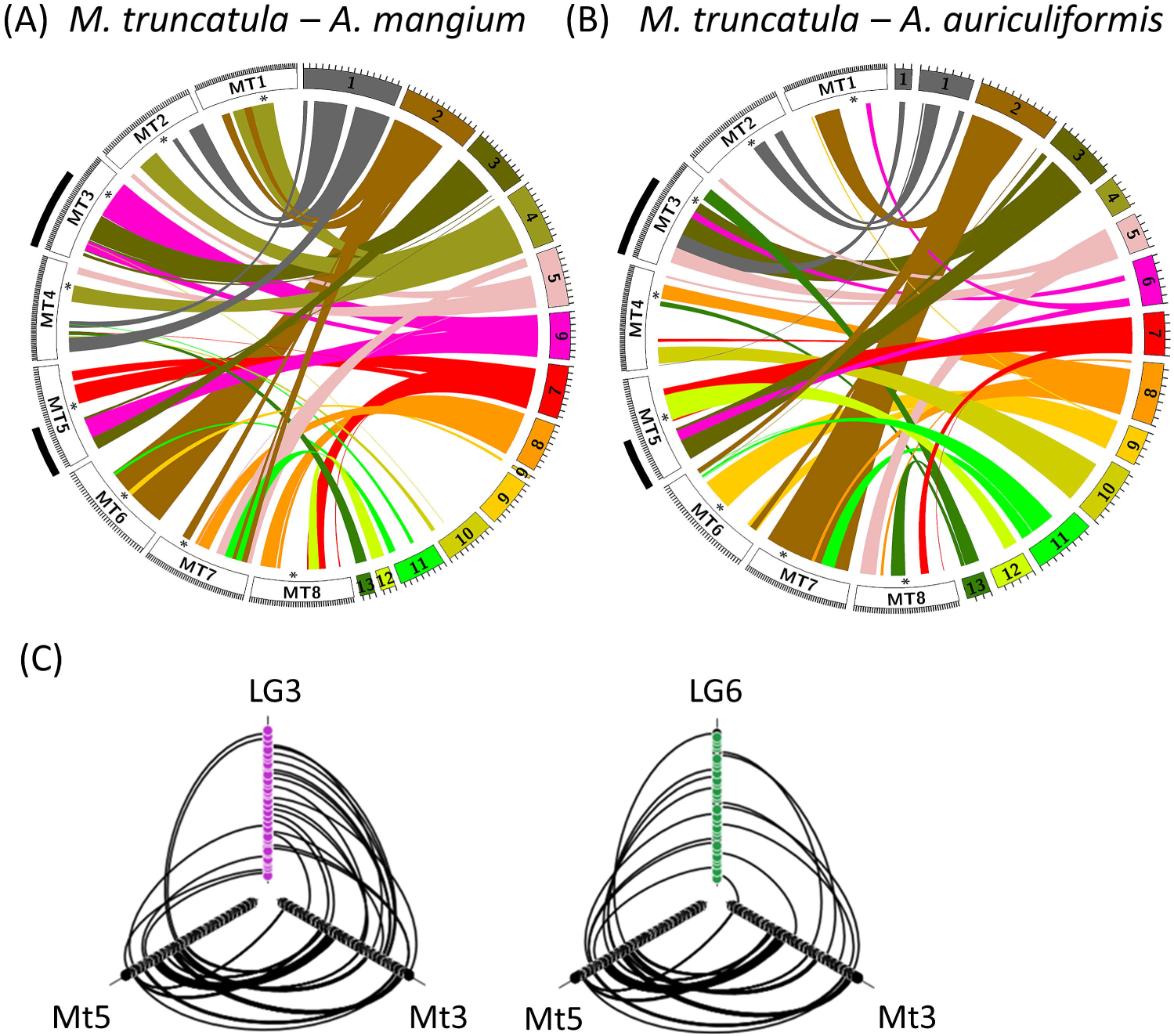
Syntenic relationship between *Acacia* linkage groups and *M. truncatula* chromosomes. Syntenic blocks detected in linkage groups of *A. mangium* (A) and *A. auriculiformis* (B) were drawn as ribbons with unique colors representing each linkage group. The asterisk represents the predicted centromere position in Mt4.0 genome. The black tiles indicate a two-to-two relationship between lower arm of *M. truncatula* chromosome 3 and 5 and LG3 and LG6 of *Acacia* genome. (C) Double Conserved Synteny of LG3 (left) and LG6 (right) from *A. mangium* with *M. truncatula* chromosome 3 (Mt3) and 5 (Mt5).

There were few overlaps between the syntenic blocks, except for LG3 and LG6 which overlapped in Mt3 and Mt5 of *M. truncatula*. A detailed examination revealed that LG3 and LG6 were syntenic to both lower arms of Mt3 and Mt5 in an interleaving pattern (Figure 1C). We observed that the genes located on one LG were homologous to two *M. truncatula* chromosomes in alternate order, a scenario also known as Double Conserved Synteny(Kellis et al., 2004). The improved *M. truncatula* Mt4.0 genome assemblies indicated that the lower arm of Mt3 is syntenic to the lower arm of Mt5, suggesting that these two chromosome arms represent the duplicated genomic blocks resulting from the Papilionoideae WGD. These findings therefore suggest that Mt3 and Mt5 are the homeologous chromosomes to *Acacia* LG3 and LG6. The observation of these two *Acacia* LGs exhibiting Double Conserved Synteny provides good evidence for a WGD in *Acacia*.

### Karyotype prediction of *M. truncatula* ancestor reveals a WGD in *Acacia*

To test the hypothesis that *Acacia* underwent a WGD like that in Papilionoideae, we predicted the Ancestral Genomic Blocks (AGBs) from *M. truncatula* modern genome (Figure 2A) and tracked their positions in the modern *Acacia* genomes based on syntenic relationships between *Acacia* LGs and the *M. truncatula* modern genome (Figure 2B). We assumed that the preduplication ancestral genome of *M. truncatula* is highly similar to that of the last common ancestor between Papilionoideae and Mimosoideae. In addition, we also used the *Vitis vinifera* modern genome as outgroup to support potential duplicated AGBs within the *M. truncatula* modern genome. Assuming all AGBs remained intact after divergence, we hypothesized that we should find two genomic regions for each AGB in each *Acacia* linkage map if the *Acacia* genome is derived from a WGD event. Alternatively, if no WGD had occurred in *Acacia*, we would find one genomic region for each AGB.

**Figure 2.**
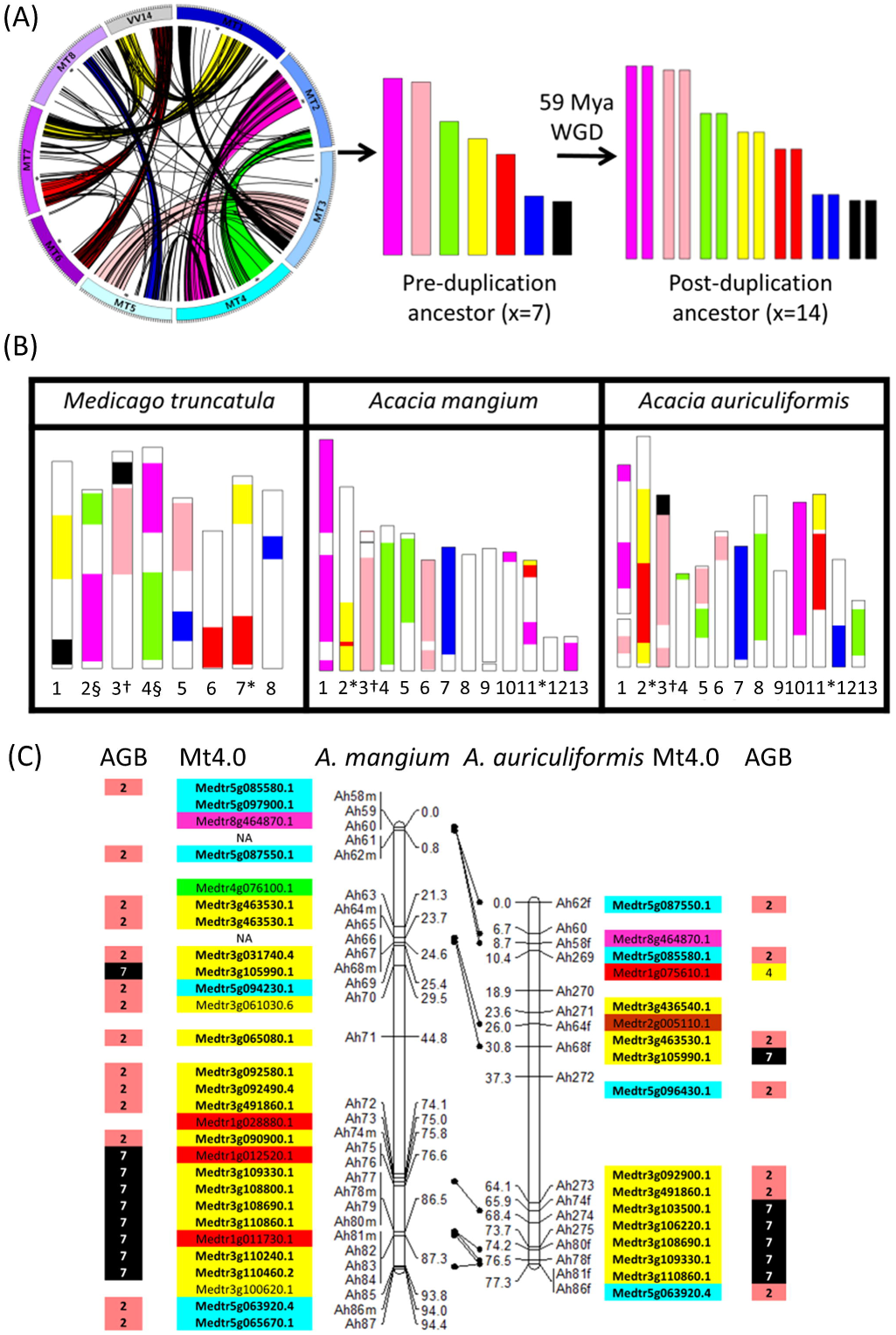
Karyotype evolution in Fabaceae. (A) Circos plot indicated the syntenic relationship within *M. truncatula* genome and between *M. truncatula* chromosome 1 (Mt1) and *V. vinifera* chromosome 14 (Vv14) which allows the identification of seven Ancestral Genomic Blocks (AGBs) as represented by karyotypes of different colour. The *M. truncatula* ancestor is predicted to have 7 and 14 pairs of chromosomes before and after 59 Mya Whole Genome Duplication (WGD). (B) The tracking of AGBs in modern genomes of *M. truncatula, A. mangium* and *A. auriculiformis*. §Mt2 and M4 both contain AGB1 and AGB3. *Mt7, LG2 and LG11 contain AGB4 and AGB5. †Mt3 and LG3 are fused product of AGB2 and AGB7. (C) Improvement of comparative genomic analysis using information of ancestral genomic blocks demonstrated in linkage group 3 (LG3) of *A. mangium* (left) and *A. auriculiformis* (right). The *M. truncatula* orthologous genes and predicted AGB membership are displayed next to the markers on LG3. One-to-one syntenic relationship showed that both LGs contain several syntenic blocks with small gene insertions, however, AGB prediction showed that both LGs contain the same AGB content and structure as the lower arm of *M. truncatula* chromosome 3.

Using this approach, we identified seven AGBs within the *M. truncatula* genome, based on 180 syntenic blocks consisting of 1,295 gene pairs within the *M. truncatula* genome, and 66 syntenic blocks based on 809 gene pairs between the *M. truncatula* genome and chromosome 14 of *V. vinifera* (Figure 2A). These AGBs consisted of five major and two minor blocks, about 11.46-20.12 Mb and 6.08-6.70 Mb long, respectively, and representing 24% of the *M. truncatula* modern genome. In the *M. truncatula* modern genome, all chromosomes have two different AGBs except Mt6 and Mt8. We failed to detect any AGB in the lower arms of Mt6 and upper arm of Mt8, which is potentially due to degraded synteny. Interestingly, Mt2 and Mt4 both contained AGB1 and AGB3, however, AGB1 was inverted in one of these two chromosomes. Since the sequenced *M. truncatula* accession is known to contain a translocation between lower arm of Mt4 and Mt8 (Kamphuis et al., 2007; Tang et al., 2014), AGB1 was most likely not found in Mt4 of other *M. truncatula* accessions. In the *Acacia* modern genomes, we observed two copies of all major AGBs, whereas all *Acacia* LGs with a few exceptions contained one AGB. We interpret the detection of only one copy of AGB7 as resulting from a possible deletion of the duplicated copy in *Acacia* genomes. In the case of LG2 and LG11, AGB5 (red) was found next to or nested within AGB4 (yellow). This was also the case in Mt7 and Vv14. The presence of two different AGBs in *A. auriculiformis* LG1 and LG5 and *A. mangium* LG11 suggests possible fusion of two different AGBs in a single chromosome, however, the evidence is rather weak as these fusions were observed in only one of the two *Acacia* linkage maps. Nonetheless, our synteny analysis supported our hypothesis that WGD has occurred in *Acacia*.

With this information, we re-evaluated the *Acacia* comparative genomic analysis using LG3 as an example (Figure 2C). In *Acacia* LG3, one-to-one comparison with *M. truncatula* genome revealed that genes from LG3 are orthologous to genes from Mt3 and Mt5 in an interleaved pattern. At first glance, one may conclude that LG3 was the fusion product of two small fragments from Mt5 at both ends of a large fragment from Mt3. In addition, the presence of genes orthologous to other chromosomes in LG3 suggested possible gene insertion from other chromosomes especially Mt1. However, the AGB prediction indicated LG3 has the same AGB content and an organization similar to the lower arm of Mt3 (Figure 2C). In both LGs, AGB7 was located at the end of AGB2 and the centromere was expected to fall at the boundary of these two AGBs. Detailed examination of blast results revealed that majority of the genes (~58%) are orthologous to Mt3 genes based on best blast hit while the rest were either absent in *M. truncatula* genome or contains hits to duplicated genes from other chromosomes because its Mt3 homologous gene is either the second best hit or missing in the genome. Based on these findings, we predicted that LG3 was the “orthologous” chromosome of Mt3. We tried to verify further that LG3 is indeed the “orthologous” chromosome of Mt3 and LG6 is the “orthologous” chromosome of Mt5 by measuring the Ks values between the two linkage groups and the two *M.truncatula* chromosomes. Our hypothesis is that if LG3 is indeed the “orthologous” chromosome of Mt3, we should observed lower ks values between markers on LG3 and Mt3 compared to ks values between markers on LG3 and Mt5. The same applied to the relationship between LG6 and Mt5. Unfortunately, the orthologous *M. truncatula* genes of all markers except one in each LG don’t have syntelogs in either Mt3 or Mt6 (Supplementary Table 2).

### Prediction of Whole Genome Duplication (WGD) event in three subfamilies

To further validate the WGD in *Acacia*, we examined the age distribution of duplicated genes using Ks analysis based on transcriptome data. This allowed us to also address other questions including: 1) Is the WGD in *Acacia* the same as in Papilionoideae? ;2) Did WGD occur in other members of Mimosoideae and Caesalpinioideae?; 3) Did members of Mimosoideae and Caesalpinioideae have additional WGD events? To look for the WGD events in Mimosoideae and Caesalpinioideae, we performed Ks analysis in nine diverse taxa consisting of three taxa representing each subfamily, namely *M. truncatula, L. japonicus* and *G. max* (Papilionoideae), *Eperua falcata, Cercis gigantea* and *C. fasciculata* (Caesalpinioideae), *P. alba, A. auriculiformis* and *A. mangium* (Mimosoideae).

The ks analysis showed clear evidence of one WGD event in both *A. auriculiformis* and *A. mangium*. Our results indicated that all nine analyzed legumes have a Ks peak ranging from mean Ks values of 0.35 to 0.68 (Figure 3A). SiZer analysis indicated that all these peaks are significant except for the peak in *P. alba*. The occurrence of many recent duplications as showed by the near zero peak has a masking effect on WGD peak of *P. alba*. We identified the Papilionoideae WGD as peaks ranging from 0.53 to 0.68 in the Ks distributions of *M. truncatula, G. max* and *L. japonicus*. The presence of a Ks peak within this range in *A. auriculiformis* and *A. mangium* not only supports the *Acacia* WGD hypothesis, but also suggests they potentially share the Papilionoideae WGD due to similar WGD age. Mean Ks values of 0.35 and 0.50 detected in the Caesalpinioideae *C. gigantea* and *E. falcata* respectively were outside the Papilionoideae WGD range. We predicted that these two species fall outside the expected Ks range maybe due to variation in synonymous substitution rates in the different lineages. No significant peak representing more recent WGD in any of the legumes analyzed was found except a peak representing 13 Mya lineage-specific WGD in *G. max*. In addition to validating new WGDs in the genus *Acacia* and *Eperua*, our results are consistent with the reported WGD ages in Cannon et al. 2015

**Figure 3.**
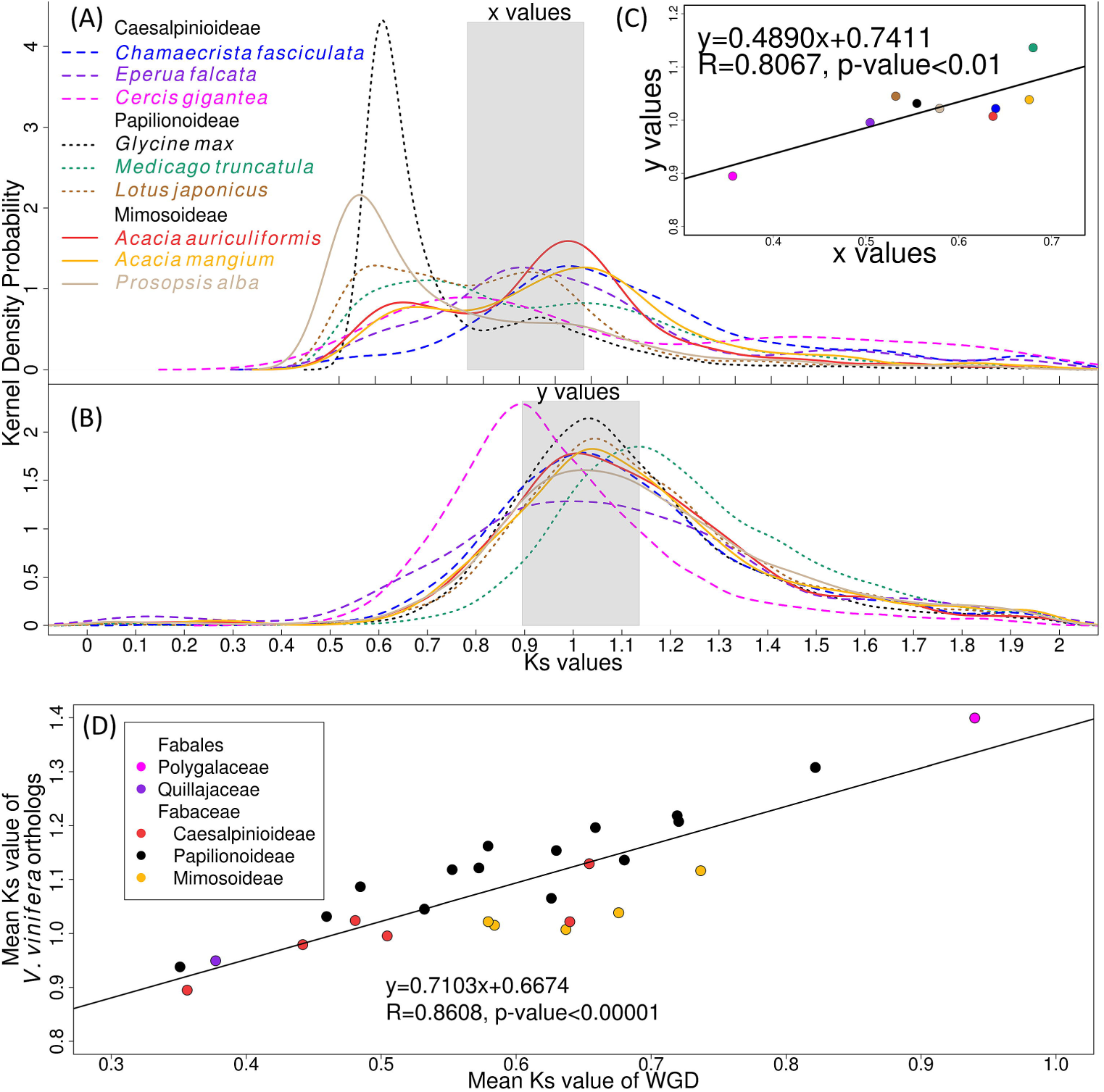
Synonymous substitution rate (Ks) estimation in Fabaceae. (A) Ks estimation for paralogs within species was used to detect Whole Genome Duplication (WGD) indicated as peaks. The mean Ks values (x values) for WGD are indicated for three subfamilies. (B) Ks estimation for *Vitis vinifera* orthologs is used to determine variation of substitution rates among the lineages. The mean Ks values (y values) for WGD are indicated for three subfamilies. (C) Linear regression between mean Ks values of *V. vinifera* orthologs (y values from figure 3(B)) and mean Ks values associated with hypothesized WGDs (x values from figure 3(A)). (D) Relationship between mean Ks values of *V. vinifera* orthologs and mean Ks values associated with hypothesized WGDs for 27 species within family Fabaceae and order Fabales. High Pearson Correlation Coefficient (R=0.8606) suggests that lineage-specific Ks variation is responsible for the variation of WGD Ks values.

To investigate whether variation of synonymous substitution rates in the legume lineages can account for the range of WGD Ks values inferred, we estimated the substitution rates for 457-4,707 orthologs between nine analyzed legumes and *V. vinifera* (Figure 3B). *V. vinifera*, which is sister to the rosids, was used commonly as outgroup to eudicots including the legumes as its genome has not undergone additional WGD and most of its chromosomes remain intact since the ancient hexaploidy event shared by all eudicots (Jaillon et al., 2007). If substitution rate was constant in all lineages, and ruling out stochastic rate variation due to ortholog sampling size, we would expect all analyzed legumes to show the same mean Ks values because they should share the same divergence time from *V. vinifera*. Our results showed that *C. gigantea* had the lowest mean Ks values (0.895) and *M. truncatula* had the highest mean Ks values (1.136) consistent with their WGD ages. We discovered that the mean Ks values of *V. vinifera* orthologs is positively correlated (Pearson correlation coefficient, R=0.8067, p-value < 0.01) with the observed Ks values for WGD despite the low sample size (Figure 3C). We improved the correlation test by using an additional 18 transcriptome datasets from 5 mimosoids, 6 caesalpinioids, 14 papilionoids, one species each from Polygalaceae and Quillajaceae which were used in Cannon et al. 2015. We found a stronger positive correlation (Pearson correlation coefficient, R=0.8606, p-value < 0.00001) between the two variables (Figure 3D). The high coefficient of determination (r^2^ = 0.74) indicated that 74% variation in mean Ks values of WGD-associated peaks can be explained by variation of synonymous substitution rates as measured by divergence age of *V. vinifera* orthologs. This result suggests that the age of WGDs of the family Fabaceae should be the same if the variation of Ks is corrected.

If WGDs in all legumes occur at the same time, are they a single WGD event shared by all legumes or multiple independent WGDs across subfamilies as proposed by Cannon et al. 2015? It is challenging to answer this question by using absolute dating of WGDs alone because the WGD ages are very similar. In our opinion, the best approach is to perform karyotype evolution and study the synteny conservation between members of Papilionoideae and Mimosoideae and/or Caesalpinioideae. To this end, we revised our karyotype evolution model which identified six ancestral blocks in modern *M. truncatula* genome and one unknown ancestral block (Supplementary Figure 2). We proposed that AGB 4 is a middle fragment of AGB 5 and they are inherited as a single chromosome as observed in Acacia. Interestingly, we observed that AGB 2 and AGB 7 are fused together as in Mt3 and Acacia LG3 suggesting that they are inherited as a single unit in legumes. We examined the positions of these two AGBs in modern genome of *V. vinifera* and were able to track the positions of AGB 2 to the start arm of Vv18 and AGB 7 in two parts, namely a part of Vv4 and a bottom part of vv18. Assuming *V. vinifera* represents the ancestral karyotype state, this observation suggests that fusion occurs after divergence from *V. vinifera* and before he divergence of Papilionoideae and Mimosoideae.

Another commonly used approach to study WGD is to construct gene trees and examine the gene duplication patterns as performed by Cannon et al. 2015. Since we have additional transcriptome datasets from a few taxa not used in Cannon et al. 2015, we are interested to re-examine six gene trees from Cannon et al. 2015 to assess whether increased taxon sampling changes gene duplication patterns and support for multiple WGD. First, we filtered gene trees which are not suitable to study WGD based on several criteria: 1) The gene is not a single copy gene; 2) Ideally, there should be two duplicated copies for each taxa with near full length sequence; 3) These genes must be clustered correctly in the same gene family; 4) Since we are only interested to study gene duplication for the WGD dated around 59Mya, this gene tree should not contain genes are from more recent or more ancient duplication events (Ks values >=2). We observed that gene tree construction using nucleotide sequences gives better bootstrap values than protein sequences. At least two of the six gene trees did not fit the above criteria because one gene tree contains only one copy from Papilionoideae and the genes from another gene tree did not belong to the same gene family (Supplementary Figure 2). The remaining four gene trees provided weak support (bootstrap value about 50) and identified Papilionoideae as monophyletic. In addition, only 2-3 taxa from Mimosoideae and Caesalpinioideae contain two duplicated copies in these four gene trees and the duplicated copies were identified as a result of very old duplication (Ks values >=2) in two gene trees. Although the remaining gene trees identified a gene duplication in Papilionoideae, we found no or little evidence of a gene duplication dated around 59Mya in Mimosoideae and Caesalpinioideae possibly due to incomplete transcriptome sequencing, gene loss and limited number of gene trees examined.

## DISCUSSION

### Karyotype evolution of modern legumes

Due to the lack of genome sequences, gene-based linkage maps have been widely used in studies involving teleost fishes and plants to infer WGD and track the distribution of ancestral chromosomes in extant genomes (Naruse et al., 2004; Schranz et al., 2006; Abrouk et al., 2010; Bordat et al., 2011; Wang et al., 2011; Guyomard et al., 2012). We constructed the first sequence-based linkage maps for *Acacia* and the second *A. mangium* linkage map after Butcher and Moran 2000 (Butcher and Moran, 2000). Although synteny was observed between *Acacia* and *M. truncatula*, our results showed that the syntenic relationship between the two genomes could not be elucidated through a simple one-to-one relationship as found in comparative genomic analyses within Papilionoideae (Sato et al., 2008; Bordat et al., 2011; Lucas et al., 2011; Varshney et al., 2011; Young et al., 2011; Cruz-Izquierdo et al., 2012; Isobe et al., 2012; Young and Bharti, 2012; Sharpe et al., 2013; Varshney et al., 2013). Direct genome comparison between Mimosoideae and Papilionoideae without local synteny data can lead to inaccurate data interpretation due to problems in assigning true orthologous genes caused by Double Conserved Synteny. Using Acacia LG3 as an example, we demonstrated that prediction of ancestral genomic blocks can assist in the identification of homeologous chromosomes and provide an important framework for comparative genomic analysis between two highly divergent genomes. Our study provides important resources for future genomics and genome sequencing projects in Mimosoideae and Caesalpinioideae.

The prediction of *M. truncatula* ancestral genome and identification of structural conservation of ancestral blocks between modern Papilionoideae and Mimosoideae genomes has provided new insights into karyotype evolution in Fabaceae. We attempted to identify seven AGBs using the M. truncatula genome Mt4.0 based on the assumption that the legume ancestor has 7 chromosomes as predicted by previous karyotyping study (Goldblatt 1981). We found two of them (AGB4 and AGB5) are linked in *V. vinifera, M. truncatula* and Acacia genomes. This is consistent with previous studies in *Lens culinaris, Vicia* sp. and *Pisum sativa* (Gujaria-Verma et al., 2014; Webb et al., 2016; Bordat et al., 2011) which indicated AGB 5 is located in the centromeric region of AGB4 in one chromosome. It is clear these two AGBs must inherited as a single chromosome in the legume ancestor and possibly eudicot ancestor. Similarly, AGB2 is linked to AGB7 in both Mt3 and Acacia LG3 but not observed in *V. vinifera*. The fusion of AGB2 and AGB7 is predicted to occur after WGD in the branches leading to modern legumes. Interestingly, the start arm of Mt3 and end arm of Mt6 are not syntenic to any region within the *M. truncatula* genome. These two regions are known to NBS-LRR megaclusters and recent study suggested that these disease resistance genes expanded through species-specific duplications (Zhong et al., 2015). In the end, our revised ancestral karyotype model predicted a total of six AGBs. It is possible that a seventh AGB is present in *M. truncatula* genome but not detected in the current genome assembly. One possible region is the start arm of Mt8 and the middle part of Mt7 which share a small number of syntenic blocks.

Our results suggested that the modern Acacia genome has not undergone major chromosomal rearrangement and most duplicated AGBs appeared to be inherited as single chromosomes after WGD. This finding is presumably true for other members of Mimosoideae and Caesalpinioideae which share similar chromosome number (n=13-14). On the contrary, most *M. truncatula* chromosomes consist of two different duplicated AGBs fused and genomic region where two AGBs fused often evolved to become the centromeric region as evident in Mt1,Mt2, Mt5 and Mt7. This suggests that end-to-end fusions led to a reduction of chromosome number in Papilionoideae (n=7-11) (Soltis and Soltis, 2012). This finding is supported by the prediction of 25 fusions involved in the formation of the modern pea genome from eudicot ancestor (Bordat et al., 2011). We demonstrated that prediction of ancestral genomic blocks, in combination with direct genome comparison, to be very useful for plants without a reference genome.

### Effect of Ks variation across Fabales on estimation of WGD age

Ks values are well-known to vary considerably across lineages and over time (Tang et al., 2008; Tuskan et al., 2006). It is thought that Ks variation among genes has strong correlation with gene expression and higher Ks is observed in annuals compared to perennial plants (Gaut et al., 2011). However, our findings suggested that Ks varition is subfamily specific and is not related to life history. This is consistent with Lavin et al. 2005 which reported fastest and slowest matK and rbcL substitution rates for Papilionoideae and Mimosoideae respectively. This huge variation of synonymous substitution rate (Ks) in legumes has several important implications on WGD inference of legumes.

Firstly, Ks variations affect the timing of WGD. If Ks variation is corrected, it would imply that all WGD within Fabales occurs at the same time which is expected to be around the same time as the Papilionoideae WGD. In Cannon et al. 2015 study, the variable WGD ages were not adjusted supporting the multiple origins of WGD in legumes. Secondly, variable Ks values can result in over-/under-estimating the divergence age between any two legume species. For example, unadjusted Ks values between a slow evolving legume (i.e. Ks=0.356 in *C. gigantea*) and a fast evolving legume (i.e. Ks=0.68 in *M. truncatula*) will lead to biased inference that the WGD in the *C. gigantea* occurred before the divergence of two species (Ks=0.574) and the WGD in *M. truncatula.* Several studies (Paterson et al., 2012; Vanneste et al., 2014) have indicated that synonymous substitution rate variation can influence the estimation of WGD age within species of the same family and suggest that absolute dating without correction for rate differences can produce misleading results.

Thirdly, Ks variation affects gene tree topology and gene duplication patterns since the branching pattern and length is dependent on mutation rate which is directly affected by synonymous rate. The gene tree approach to investigate the origin of duplication on large scale gene family is a commonly used method. This method assumes that if a gene is duplicated, one copy of the gene branches off at the origin of gene duplication and further undergone speciation to form a monophyletic group consisting of orthologous genes from closely related species. The location of WGD can be inferred by examining the location of the gene duplication origin in the gene tree when large number of species are used. Further investigation is required to study how Ks variation affects gene tree topology which can lead to biased inference on WGD. There is a need to re-examine gene duplication patterns in all legumes using a more robust approach such as phylogenomic methods which corrects for Ks variation.

### A single WGD hypothesis in all legumes

In this study, we argue that our Ks analysis, along with the synteny patterns observed, all provide evidence for a single WGD hypothesis for a number of reasons. Firstly, parsimony reasoning would favour a single WGD event hypothesis over one involving multiple ‘ad hoc hypotheses’ as these are more costly. For instance, the success of polyploids depends on their abilities to adapt in new environments, compete with diploid progenitors, producing fully fertile progenies and maintaining genome stability (Comai, 2005). Secondly, the interval between origin and diversification of Fabaceae was estimated to be about 1-2.5 million years (Lavin et al., 2005) and therefore, having multiple WGD within a short time frame is unparsimony. The parsimony theory was similarly applied to WGD in soybean and *M. truncatula* when WGDs were first discovered (Pfeil et al., 2005) despite the fact that both species show different WGD time and it was proven to be true by subsequent studies.

The karyotype evolution in Papilionoideae and Mimosoideae is less complex compared to Arabidopsis spp., grasses and mammals which consists of several types of chromosomal rearrangement such as reciprocal translocations, fusions and inversions (Yogeeswaran et al., 2005; Lysak et al., 2006; Murat et al., 2010). The observation that modern Acacia genomes have undergone relatively few chromosomal rearrangements and more closely resemble the post-duplication ancestral genome of *M. truncatula* in terms of chromosome number and AGB organization, compared to the modern *M. truncatula* genome provided strong evidence that a WGD predates the divergence of Papilionoideae and Mimosoideae. If WGDs occurred separately in Papilionoideae and Mimosoideae, we should expect that the modern genome of Mimosoideae modern genome would accumulate more structural changes. Another interesting observation is the fusion of AGB2 and AGB7 found in modern genome of *M. truncatula* and *Acacia* which can potentially shred some lights in the phylogeny placement of this WGD event. Parsimony reasoning would favour this fusion occurring before the divergence of Papilionoideae and Mimosoideae and after a single WGD over the alternative hypothesis that this fusion occurs separately for *M. truncatula* and *Acacia* after separate WGD (Supplementary Figure 2). The main limitation in pinpointing the exact location of WGD using synteny approach is the limited number of markers in the *Acacia* linkage maps. The linkage map needs to be improved to contain sufficient markers to identify syntelogs which are sets of genes derived from the same ancestral genomic region.

Our findings suggest that a single WGD event near the base of Fabales was associated with the initial diversification of the Fabaceae and subsequent chromosome number reduction and high synonymous substitution rates contributed to the success and diversification of Papilionoideae. Since WGDs of the same age are also found in Polygalaceae and Quillajaceae, the phylogenetic placement of this WGD event is predicted to be near the base of Fabales. This timing is within Cretaceous–Tertiary boundary (65 Mya) where many independent WGD occurred in plant families and consistent with absolute dating of Papillionoideae WGD (Fawcett et al. 2009). This single WGD hypothesis is consistent with the observation of significant time lag between WGD and radiation has been observed in Brassicaceae, Poaceae, Fabaceae, Asteraceae, Solanaceae, and angiosperms (Schranz et al. 2012). We think that polyploidy may have increased the survival chances and recolonization capacity of plant lineages during and/or after the KT mass extinction.

## METHODS

### *Acacia* linkage map construction and comparative analysis with *M. truncatula*

A total of 768 gene-based Single Nucleotide Polymorphism (SNP) molecular markers were used for linkage map construction. The development, genotyping and selection of these markers employed for the mapping pedigree in this study have been described previously in other studies (Wong et al., 2012; Wong, 2013). Linkage mapping was performed in Joinmap 4.0 (Van Ooijen, 2006) using default settings and the Kosambi mapping function. Independent linkage map were constructed for each parent based on 123 individuals. Segregation distortion (p≤0.05) was checked using the Chisquare test and the accuracy of the genotype data was improved by identifying significant double crossover events (p<0.05). Markers were grouped at ≥ LOD 4 with linkage maps visualized using Mapchart 2.2 (Voorrips, 2002). Comparative genomic analysis between *Acacia* and *M. truncatula* was carried out independently using each parental linkage maps. Marker sequences were re-aligned against *M. truncatula* CDS version Mt4.0v1. The best ortholog for each *Acacia* sequence was predicted based on the highest total score of the alignments. Due to large divergence time between *Acacia* and *M. truncatula*, syntenic blocks were predicted using OrthoclusterDB (Ng et al., 2009) with somewhat relaxed parameters (option –r –i 10 –ip 50 –o 10 –op 50) which requires two syntenic blocks to share at least two orthologous genes, a maximum of 10 intervening genes and 50% gaps in a syntenic block regardless of its strandedness. The syntenic blocks were manually inspected by removing syntenic blocks with only two markers and large gap. The syntenic relationships between *Acacia* and *M. truncatula* genome were visualized using Circos version 0.65 (Krzywinski et al., 2009). To visualize Double Conserved Synteny in *Acacia* LG3 and LG6, syntenic relationships between *Acacia* and *M. truncatula* genome and syntenic relationship between Mt3 and Mt5 were used to plot a hiveplot using Jhive-0.2.7 (Krzywinski et al., 2012).

### Karyotype evolution in Fabaceae

To determine whether WGD has occurred in *Acacia* using a synteny-based approach, we predicted the Ancestral Genomic Blocks (AGBs) from pre-duplication ancestor of *M. truncatula* genome by examining the syntenic relationships within *M. truncatula* genome. Synmap (http://genomevolution.org/CoGe/SynMap.pl) was used to identify syntenic blocks within the *M. truncatula* version Mt4.0 genome by generating a dotplot of the CDS against itself using default parameters. Synmap is a Perl pipeline from CoGe which implements QUOTA-ALIGN algorithm and identifies gene pairs using Blast and DAGchainer (Lyons et al., 2008; Haas et al., 2004; Lyons et al., 2008; Tang et al., 2011). The DAGchainer output file containing syntenic blocks was downloaded. In addition, syntenic relationships between *V. vinifera* and *M. truncatula* genome were used to predict an AGB in Mt6 and 7 using the same approach. These syntenic relationships were visualized using Circos version 0.65. Major and minor AGBs of at least 10 MB and 5 MB respectively were identified by visual inspection of the Circos plot.

To track the positions of these AGBs in linkage maps of *Acacia*, we used two methods to infer the conserved synteny between the AGBs and *Acacia* genome. In the first method, each AGB was assumed to be a part of the *M. truncatula* ancestral genome and was represented by two genomic regions in *M. truncatula* modern genome. We made the assumption that an *Acacia-Medicago* syntenic block is syntenic to an AGB if it overlaps or falls within one of the two genomic regions of that AGB. In addition, an *Acacia-Medicago* syntenic block which is syntenic to an AGB must contain at least two orthologous genes within the two genomic regions of that AGB. Using this information, we plotted the distribution of AGBs in *M. truncatula* genome, *A. mangium* and *A. auriculiformis* LGs using R. In the second method, we examined whether a gene in the LGs is orthologous to any gene within an AGB. We re-evaluated the comparative analysis of *Acacia* LG3 by comparing this AGB membership method and the initial direct *M. truncatula* genome comparison method. Refer Supplementary Table 1 online for detailed information about linkage maps and ancestral genomic blocks.

### Prediction of Whole Genome Duplication (WGD) events in Fabaceae

Synonymous substitution rates (Ks) were estimated from genes or transcriptome datasets using modified methods from Blanc and Wolfe 2004 (Blanc and Wolfe, 2004) for *V. vinifera* as an outgroup and three members of each Fabaceae subfamily, namely *M. truncatula, L. japonicus* and *G. max* (Papilionoideae), *E. falcata, C. gigantea* and *C. fasciculata* (Caesalpinioideae), *P. alba, A. auriculiformis* and *A. mangium* (Mimosoideae), and additional 18 species across family Fabaceae and order Fabales from Onekp project used in Cannon et al. 2015 (Cannon et al., 2015). Pairwise Ks values were estimated for paralogous genes within each dataset and orthologous genes between any two datasets using a modified Python script from Tang et al. 2010 (Tang et al., 2010), which: 1) aligned pairwise protein sequences using ClustalW2 (Larkin et al., 2007);2) aligned nucleotide sequences based on protein alignments using PAL2NAL v14 (Suyama et al., 2006); and, 3) estimated Ks values using CODEML (default settings except runmode −2) in PAML 4.7 (Yang, 2007). To identify paralogous genes within a dataset, all-against-all alignments were performed using Blast+2.2.27 Blastn megablast algorithm. Any alignment with less than 40% identity, 300 bp length and 70% length of the shortest sequence was removed. For the transcriptome datasets, prior to Ks estimation, nucleotide and peptide sequences were predicted from aligned regions using a custom Biopython script. Any paralogous pairs with Ks values equal to zero which result from local duplications or alternative splicing were removed. Single linkage clustering and calculation of median Ks for each cluster were performed based on methods from Blanc and Wolfe (Blanc and Wolfe, 2004) using a custom Python script. The same method was used to calculate the Ks values for orthologous genes between any two datasets except the orthologous sequences must be the best hit for each other based on total alignment scores. All distributions of the Ks values were plotted using kernel density estimation in R and SiZer (Chaudhuri and Marron, 1999) was used to look for significant peaks in a Ks distribution plot. The mean Ks values of the Ks distribution were determined in R based on the highest values within a density function. To determine the relationship between substitution rate variation and age of WGD, the mean Ks values of *V. vinifera* was plotted against mean Ks values of WGD and Pearson’s correlation coefficient and linear regression were calculated using R. The full details of all transcriptome datasets and Ks values are available in Supplementary Table 2 online.

To evaluate whether the gene duplication patterns support single or multiple WGDs in legumes, we studied six gene trees with full tree topology published in Cannon et al. 2015, namely Figure 4A, Figure 4B, Figure S2A, Figure S2B, Figure S2C, Figure S2D. We tried to reproduce the gene trees by using the same gene sequence from these gene trees besides adding sequences of new species from our study which are identified using Blast+2.2.27 Blastp algorithm (Camacho et al., 2009) and Orthomcl v2.0.9 (Li et al., 2003). To make sure the genes fit the assumption that they are duplicated genes from WGD around 59 Mya, we examined the genes in all six gene trees by checking Orthomcl grouping on LegumeIP website (Li et al., 2012) and pairwise Ks values. Any duplicated genes from very ancient WGD (Ks values >= 2.0) or recent duplication and single copy gene were removed from further analysis. We aligned the genes (both nucleotide and protein sequences) using MAFFT and Clustal Omega from EMBL-EBI web services (Sievers et al., 2011; Li et al., 2015; Katoh and Standley, 2013) and constructed the gene trees using RAxML (Stamatakis, 2014) on Cipres Science Gateway (Miller et al., 2011) before visualization using Newick Viewer (Boc et al., 2012).

## SUPPLEMENTARY DATA

Supplementary Figure 1. Construction of linkage maps for *A. auriculiformis* and *A. mangium* and comparative genomic analysis with *M. truncatula* genome.

Supplementary Figure 2. Revised karyotype model including fusion event of AGB2 and AGB7 and re-examination of gene tree construction.

Supplementary Table 1. Description and table of SNPs including Blast results of SNP markers to *Medicago truncatula* genome, syntenic blocks and predicted ancestral genomic blocks.

Supplementary Table 2. Transcriptome resources and synonymous substitution rate estimation for three Fabaceae subfamilies.

## ACKNOWLEDGEMENTS

We acknowledge fundings and fellowships from Universiti Kebangsaan Malaysia and Ministry of Science, Technology and Innovation, National Science Fellowship and International Tropical Timber Organization. We are extremely thankful to University of Tasmania for hosting an attachment. We would like to thank Martin Krzywinski and Eric Schranz for constructive suggestions on graphical presentation of the manuscript and Dr. Norwati Muhammad and Dr. Mohd Zaki Abdullah of FRIM for development of the hybrid mapping populations.

## AUTHOR CONTRIBUTIONS

MW, WR, RV, JF and CH participated in the design and analysis of linkage map. MW designed the comparative genomics analysis and drafted the manuscript. MW, RV, CC and FB analyzed the comparative genomics results. WR secured funding and coordinated the project. All the authors read and approved the final manuscript.

